# Home ground advantage: selection against dispersers promotes local adaptation in wild Atlantic salmon

**DOI:** 10.1101/311258

**Authors:** Kenyon B. Mobley, Hanna Granroth-Wilding, Mikko Ellmen, Juha-Pekka Vähä, Tutku Aykanat, Susan E. Johnston, Panu Orell, Jaakko Erkinaro, Craig R. Primmer

## Abstract

A long-held, but poorly tested, assumption in natural populations is that individuals that disperse into new areas for reproduction are at a disadvantage compared to individuals that reproduce in their natal habitat, underpinning the eco-evolutionary processes of local adaptation and ecological speciation. Here, we capitalize on fine-scale population structure and natural dispersal events to compare the reproductive success of local and dispersing individuals captured on the same spawning ground in four consecutive parent-offspring cohorts of wild Atlantic salmon (*Salmo salar*). Parentage analysis conducted on adults and juvenile fish showed that local females and males had 9.6 and 2.9 times higher reproductive success than dispersers, respectively. Our results reveal how dispersal disadvantage in reproductive success may act in natural populations to drive population divergence and local adaptation over microgeographic spatial scales without clear morphological differences or physical barriers to gene flow.

## Introduction

The pattern of individuals exhibiting higher fitness in their local habitat compared to individuals originating from others, and the process leading to such a pattern is known as local adaptation (Williams 1966; Kawecki & Ebert 2004; Blanquart *et al.* 2013). Local adaptation arises when the optimal phenotype varies geographically, primarily due to environmental heterogeneity. As a result, populations may evolve locally advantageous traits under divergent selection (Williams 1966; Kawecki & Ebert 2004). Studies of local adaptation can provide insights into the evolution and maintenance of diversity, species responses to climate change, and ultimately, the processes of speciation and extinction (Schluter 2001; Turelli *et al.* 2001; Leimu & Fischer 2008; Fitzpatrick & Keller 2015). Therefore, understanding the forces such as selection, migration, mutation and genetic drift that may promote or constrain local adaptation is an important aim in biology (Adkison 1995; Kawecki & Ebert 2004; Hereford 2009; Fraser *et al.* 2011; Yeaman & Whitlock 2011; Savolainen *et al.* 2013; Fitzpatrick *et al.* 2015).

One ecological selection pressure that can limit gene flow between populations and influence local adaptation is disperser disadvantage, or selection against dispersers and their offspring, also known as immigrant inviability (Hendry 2004; Nosil *et al.* 2005). Disperser disadvantage is the result of pre- and post-zygotic reproductive barriers to dispersal including reduced survivorship and reproductive success of migrants via mortality, mate choice, assortative mating, and reduced survivorship and fitness of hybrid offspring (Nosil *et al.* 2005). Consequently, disperser disadvantage can be interpreted as a reproductive isolating mechanism that promotes local adaptation and ecological speciation (Rundle Howard & Nosil 2005).

Natural dispersal events have the potential to illuminate the importance of reproductive success in shaping local adaptation, yet few studies comparing reproductive success between local and migrating individuals are conducted in nature (but see Orell *et al.* 1999; Hansson *et al.* 2004; Peterson *et al.* 2014). Rather, reciprocal transplant experiments are commonly employed for testing whether local individuals have higher relative fitness (Nosil *et al.* 2005; Hereford 2009; Anderson *et al.* 2015). However, transplant experiments have several limitations that are rarely recognized. First, for practical reasons many reciprocal transplant experiments measure fitness-related traits, rather than directly measuring reproductive success of local and foreign pairings, yet this is a key component of the strength of selection against migrants and the cost of adaptation to different environments (Nosil *et al.* 2005; Tobler *et al.* 2009). Second, reciprocal transplant studies rarely, if ever, acknowledge the potential effects of the choice of ‘migrant’ individuals in reciprocal transplants. For example, randomly selected individuals from a population may not reflect true natural dispersers that express different behavioral or physiological phenotypes, and thus bias the results of the fitness estimation in non-local environments (Haag *et al.* 2005; Bowler & Benton 2007; Cote *et al.* 2010; Cote *et al.* 2011; Edelaar & Bolnick 2012). An alternative approach to investigate local adaptation, particularly in natural systems, is to capitalize on genetic methods to identify natural migrants and compare the fitness between local and migrant individuals.

In recent decades, salmonid fishes have become a model system for studying local adaptation in natural systems (Taylor 1991; Adkison 1995; Lu & Bernatchez 1999; Fraser *et al.* 2011; Westley *et al.* 2013; O’Toole *et al.* 2015). Many salmonids are anadromous, migrating from freshwater spawning grounds to marine habitats, and demonstrate high fidelity to their natal spawning sites on their return migration (Quinn 1993; Hendry *et al.* 2003; Bett *et al.* 2017). Atlantic salmon (*Salmo salar*) exhibit extensive variation in life history strategies related to the timing of sexual maturity and migration strategies (Quinn 1993; Hendry *et al.* 2003; Barson *et al.* 2015; Bett *et al.* 2017). A small proportion of salmon disperse to new spawning grounds upon returning from their marine migration (Fleming 1996; Garant *et al.* 2003; Fleming & Einum 2011). Here, we take advantage of natural dispersal events in a wild Atlantic salmon population complex in the Teno River system of northern Finland (Vähä *et al.* 2017; Erkinaro *et al.* 2018) (Fig. 1) to compare the reproductive success of local individuals spawning in their natal environment versus individuals that have dispersed from other populations within the complex. Using data from four consecutive parent-offspring cohorts in a large spawning ground, we found that local individuals have a distinct and consistent fitness advantage over dispersers in both males and females. Similar results were observed in a single cohort from a second spawning area. Our results imply that local selection against dispersers lay the groundwork for local adaptation and population divergence in this species.

## Materials and Methods

### Study system

The Teno River is a large river in northern Europe (68–70°N, 25–27°E) that forms the border between Finland and Norway and drains north into the Tana Fjord at the Barents Sea (Fig. 1). The river is characterized by a high level of temporally-stable genetic sub-structuring between tributaries (Vähä *et al.* 2007, 2008), and genetically distinct populations within the mainstem are also reported (Aykanat *et al.* 2015; Vähä *et al.* 2017). This study focuses on a 2km river stretch at the mouth of the Utsjoki tributary including large gravel beds suitable for spawning in the Teno river mainstream (lower Utsjoki: 69°54′28.37"N, 27°2′47.52"E, Fig. 1). Fish from the lower Utsjoki are genetically identical to individuals captured from the lower Teno mainstem (Vähä *et al.* 2017) (Fig. 1). We also sampled adults and offspring at a second spawning location, Akujoki River over one cohort year (2011-2012; Fig. 1, see Appendix S1 in Supplementary Information) but for simplicity, we only present results from lower Utsjoki in the maintext.

### Sampling

Anadromous adults were sampled in September-October on the spawning grounds, approximately 1-2 weeks prior to the commencement of spawning, and had attained full secondary sexual characteristics. Throughout the study, “cohort year” refers to the spawning year when adults were captured and offspring were fertilized, although offspring were sampled in the subsequent calendar year. In total, four parent-offspring cohorts were sampled between 2011 and 2015. All sampling was approved by local environmental agencies and water management co-operatives, and conducted according to the relevant national guidelines.

Adults were caught using gill nets except for a few males caught by angling. Fish were transported into large keep nets by boat and then sexed, weighed and measured for total length (tip of snout to end of caudal fin). Scale and fin tissue samples were taken for phenotypic and molecular analyses, respectively. We released fish back into the river after sampling. During sampling, four adults died in the net (two disperser females, one disperser male and one local male). One of the dead females (caught October 1^st^) was observed to already have released her eggs and was therefore retained as a potential parent. The other three dead individuals were excluded from all analyses.

Juveniles were sampled 10-11 months later (around 2-3 months after they are expected to have emerged from the nests in the stream bed gravel) by comprehensively electrofishing all accessible river sections on or close to the spawning areas (Fig. 1). Genetic samples were collected from all juveniles by removing a portion of the adipose fins or adipose- and anal fins. After sampling the juveniles were returned into the river.

### Sea age at maturity

Sea age at maturity, defined as the number of years an individual spent at sea before returning to spawn (measured in seawinters (SW)) was determined for adults captured on the spawning ground using scale growth readings following internationally agreed guidelines (ICES 2011) as outlined in Aykanat *et al.* (2015). For one female and 15 males from which scales could not be obtained, sea age at maturity was estimated based on their weight. Briefly, data from known-sea age fish were used to define a normal distribution of weight in each sea age class, the likelihood of the weight of each unknown-age individual occurring in each sea age class was calculated, and each of these individuals was assigned the most likely sea age class for their weight (Fig. S1 in Appendix S1).

### DNA extraction

DNA of adults and juveniles was extracted from 1-2 mm^3^ of ethanol-preserved fin tissue with the QIAamp 96 DNA QIAcube HT Kit using Qiacube HT extraction robot using the recommended tissue extraction protocol with the following modifications: washing with the AW2 buffer was conducted twice, top elute buffer was not used and samples were incubated for 5 minutes before vacuum in the elution step. The final elution volume (AE buffer) was 100 μl.

**Figure 1.**
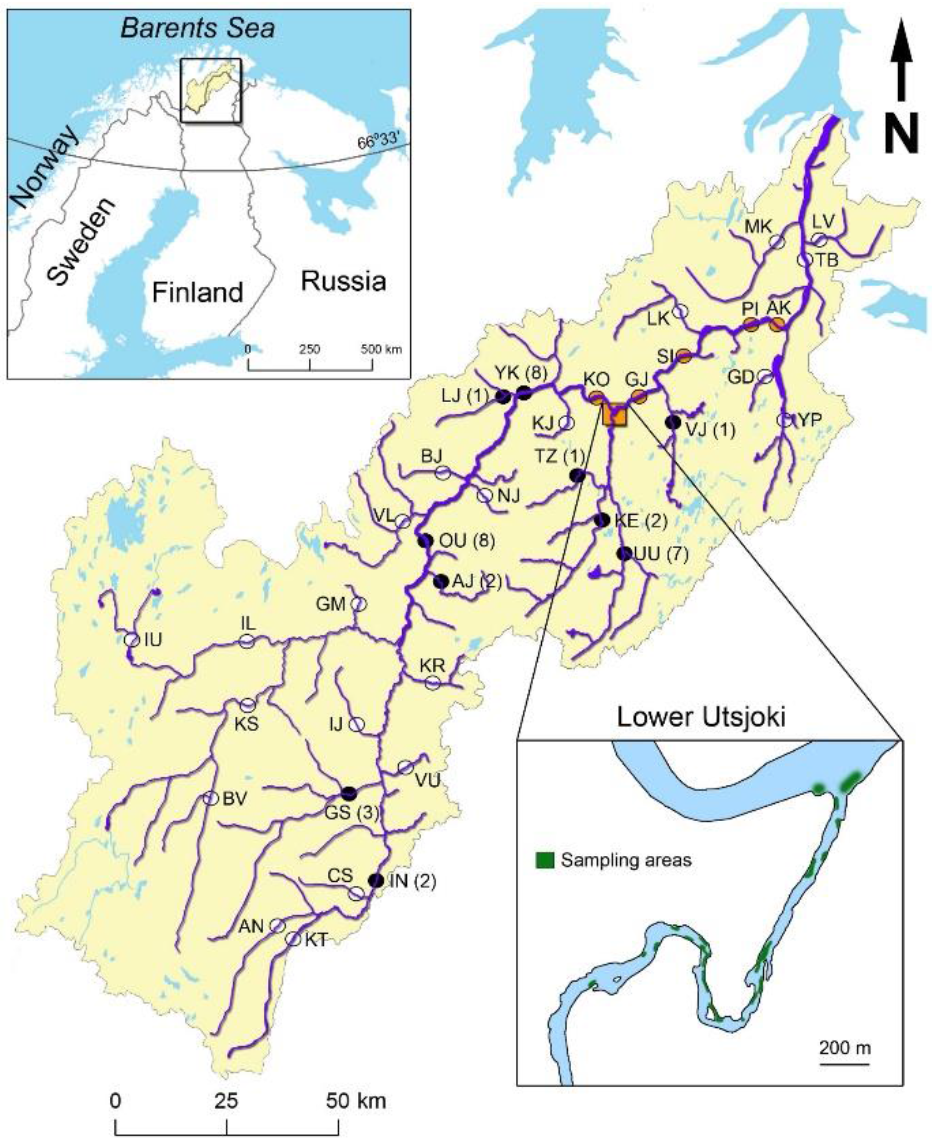
Locations sampled for baseline populations (indicated with circles) in the Teno river basin (Vähä *et al.* 2017). The orange square represents the lower Utsjoki study site and orange circles represent locations in the Teno mainstem that were considered as ‘local’. Black circles represent locations where spawning adults were assigned as dispersers with the number of assigned individuals noted in parentheses. Open circles represent baseline populations to where none of the breeding adults were assigned. Lower right inlay shows areas in green where adult and juveniles were sampled in the lower Utsjoki sampling location. Abbreviations for sampling locations can be found in supplemental data.

### Microsatellite genotyping

All adults and juveniles were genotyped using 13 microsatellite loci (panel 1 as outlined in Aykanat *et al.* (2014) excluding locus *Sssp2210*) for parentage assignment. Adults were genotyped with an additional 17 loci (30 total loci) in order to improve their assignment to their population of origin (panel 2 as outlined in Aykanat *et al.* (2014) plus *Sssp2210* from panel 1 and *MHCII* as outlined in Ozerov *et al.* (2017)). PCR amplification was performed in 6μl (panel 1) or 8 μl (panel 2) volume using Qiagen multiplex master mix (Qiagen Inc. Valencia, CA, USA) with 0.1 to 0.4 μM of each primer. The amplification was carried out according to the manufacturer’s standard protocol with annealing temperatures of 59 °C (Panel 1 MP1) or 60 °C. Visualisation of PCR products was achieved on a capillary electrophoresis-based ABI PrismTM 3130xl instrument (Applied Biosystems). Samples were prepared by pooling 1.6 μl of MP1 and 1.7 μl MP2 with 100 μl of MQ-H_2_O (panel 1) and 1.5 of MP1 and 1.5 μl of MP2 with 100 μl of MQ-H_2_O (panel 2). HiDi formamide (Applied Biosystems) and GS600LIZ size standard (Applied Biosystems) were mixed by adding 10 μl of HiDi and 0.1 μl of size standard/reaction. 10 μl of this mix was added in each well together with 2 μl of pooled PCR products. Before electrophoresis samples were denatured at 95 °C for 3 minutes. Alleles were visually inspected with Genemarker v.2.4 (SoftGenetics).

### Population assignment

Adults were assigned to their population of origin using the ONCOR program (Kalinowski *et al.* 2007; Anderson *et al.* 2008). Population baseline data for individual assignment consisted of 3323 samples originating from 36 locations as described in Vähä *et al.* (2017). Microsatellite data indicated that the baseline sampling sites extending 45km downstream and 5km upstream of the spawning location were not significantly differentiated from each other (Vähä *et al.* 2017) (Fig. 1). Therefore, microsatellite data from these five sampling sites were pooled to form one baseline population (the ‘mainstem lower’ baseline in Vähä *et al.* (2017)) and adults assigned to this baseline sample were considered local (see Table S1 in Appendix S1).

### Reproductive success

Reproductive success was quantified as the number of offspring assigned to an adult, following parentage assignment of all offspring. Pedigrees were constructed for each parent-offspring cohort separately using the package MasterBayes (Hadfield *et al.* 2006) V2.55 in the program R (R Core Team 2017). MasterBayes implements a Bayesian approach using MCMC sampling to estimate the most likely pedigree configuration while simultaneously estimating the unsampled population size, thus reducing bias in parentage assignments by allowing for uncertainty in all model parameters.

The pedigree model used fixed genotyping error rates in a two-level model (Wang 2004), calculated by re-genotyping 190 randomly chosen samples from 24 initial plate runs and comparing the two runs. Allelic drop-out (E1) was calculated as the frequency of genotypes that were homozygous in one run and heterozygous in the other, which yielded more conservative error rates than MicroDrop, a dedicated tool to estimate allelic drop-out in genetic data without repeat samples (Wang *et al.* 2012). Stochastic error rate (E2) was calculated as the frequency of alleles that were scored differently in the two runs, conservatively also including one allele from all putative cases of allelic dropout. E1 and E2 were calculated separately for each locus. Across all 13 loci, mean E1 was 0.20% and mean E2 was 0.24%.

We calculated allelic frequencies from the parental genotypes to prevent skewing by family groups produced by particularly fecund parents. Alleles from unsampled parents, present in the offspring but not in the parental genotypes, were added manually to the parental genotypes at low frequency. A simulation analysis showed that, among offspring with confidently assigned parents, this marker panel identified the true (positively identified or unsampled) mother with 96.6% accuracy and true father with 93.3% accuracy (10 pedigree runs on genotypes simulated from the final pedigree). Errors in the simulation involved unsampled parents or low-confidence assignments, with different known parents assigned in different runs in only 0.2% of dam assignments and 0.3% of sire assignments. For offspring with 10 or 11 loci typed, one mismatch with potential parents was allowed, and for those with 7-9 loci typed, no mismatches were allowed. Of 2552 offspring typed at 10 or more loci and confidently assigned at least one parent, none showed any mismatches with the assigned parent(s). Of 264 adults and 5341 offspring, 118 offspring with fewer than 7 loci successfully genotyped were excluded; all parents were successfully typed with at least 11 loci (204 at all 13 loci). Final sample sizes and assignment probabilities are shown in Tables S2 and S3 in Appendix S1.

Priors for the Bayesian inference were chosen to be broad but informative. The number of unsampled parents (unsampled population size) was estimated for both mothers and fathers in association with the pedigree estimation through MCMC sampling from the prior distribution, specified with a mean of four times the sampled population size (Aykanat *et al.* 2014), and variance calculated as 1.5 – 0.25 * sampled population size, which encompassed likely parameter space. The model was run for 70,000 iterations after a burn-in of 5,000, thinning every 2 iterations (Link & Eaton 2012). The modal pedigree configuration was extracted from the posterior distribution of pedigrees, and assignments with a likelihood of at least 90% were used in the analyses.

### Statistical modelling

Relationships between parental traits including sea age at maturity, body size, origin (local or disperser), and reproductive success were tested in a generalized linear modelling framework using a zero-inflated model from the R package pscl (Zeileis *et al.* 2008; Jackman 2017). The distributions of number of offspring contained a large proportion of zeros, which could arise if parents did not attempt to breed in the sample area but instead migrated elsewhere, and/or if parents bred but did not produce any offspring that were sampled or survived to swim up. The zero-inflated model was a two-component mixture model, accounting for zeros both in a binary term for the probability of the unobserved state (did or did not reproduce) and as zero counts as part of the proper count distribution. The zero-inflation probability was modelled in a binomial model with a logit link and the counts in a Poisson regression with a log link. The response was the number of offspring produced and all effect sizes are presented on the scale of the predictor in this log-linear model. Annual differences in sampling effort were addressed by offsetting both parts of the model by log(number of offspring sampled in that parents’ sampling year). Males and females were analyzed separately to account for expected differences in the distributions of reproductive success. In addition, variation in phenotypes (sea age at maturity and weight) between local and disperser parents was tested using Gaussian linear models in the R package lme4 (Bates *et al.* 2015).

All models of reproductive success included a main effect of sea age at maturity as older fish tend to be more successful breeders, and indeed, later-maturing fish of both sexes produced more offspring (Fig. 2, Table S4 in Appendix S1). In addition, between-year differences in the number of adults and juveniles sampled could affect reproductive success measures by influencing the likelihood that parents and offspring are identified in the pedigree. Hence, the number of offspring and adults of the relevant sex sampled each year (annual sample size) was included in all models. In models including these background variables, we tested for a main effect of individual origin (local or disperser). To further examine the role of sea age at maturity, the interaction between individual origin and sesa age was tested, using sea age at maturity as a two-level factor (termed “age class”; 1 and 2SW vs. 3SW or older in females, 1SW vs. 2SW or older in males) to account for the low numbers of older disperser fish. All these effects were tested only in the count component of the model. In all models, zero-inflation was addressed using a constant (intercept-only) binomial component of the mixture model.

To assess between-year consistency in the effect of origin, the modelling of origin as a main effect was repeated with each sampling year removed in turn, using a dataset that combined males and females, to allow for very small annual sample sizes of females and/or disperser fish. This model was the same as the main model except that it included an interaction between origin and sex, to allow for differences in effect size between males and females, and annual sample size was taken as the total number of adults (rather than of each sex) sampled each year.

We investigated local and disperser mating patterns by examining the mating success and whether breeding was assortative according to origin. We calculated mating success as the number of unique mates for each parent identified through parentage analysis. We examined mate choice among the 55 pairs where both parents were positively identified; for each individual in these pairs, the proportion of mates who were local and the mean weight of mates were calculated. The relationship of all responses with individual origin was tested in the R package nlme (Pinheiro *et al.* 2017) V3.1-137 using a Poisson generalized linear model (GLM) for the number of mates, a Poisson mixed model (GLMM) for the number of offspring per pair with mother ID fitted as a random effect to account for many females being represented more than one pair, a Gaussian model including a main effect of mate sea age for mate weight, and a quasibinomial GLM for the proportion of local mates. In the latter two, the response was weighted by the number of mates.

**Figure 2.**
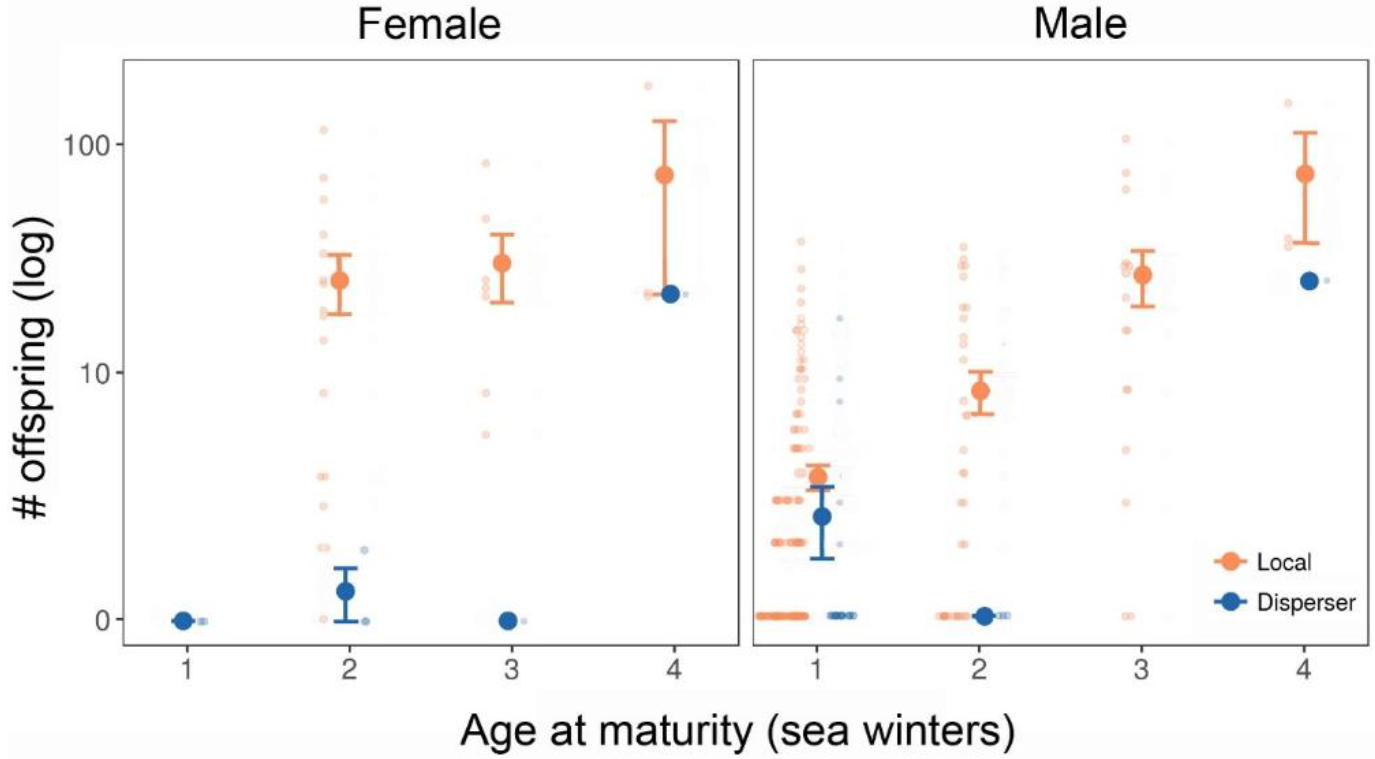
The effect of origin (local or disperser) and sea age of maturity (sea winters) on reproductive success. Large circles with error bars indicate the mean and ± standard error of the mean, small circles show individual data points. For clarity, points are jittered on the x axis. No very young (1SW) local females and only one old (3-4SW) disperser male were recorded.

## Results

### Population assignment

A total of 264 adults (230 males and 34 females) and 5223 juveniles (all < one year old) were collected in the lower Utsjoki location of the Teno River over four consecutive parent-offspring cohort years from 2011-2015 (Table S2 in Appendix S1). There was a significant 7:1 male-bias in the sex ratio (two-sided X^2^ = 55.06, P < 0.0001) that was consistent over the four cohort years.

Genetic population assignment using a conditional maximum likelihood approach based on 30 microsatellite markers indicated that 231 (87.5%) of the adults (88.7% of males and 79.4% of females) originated from the same genetic population near the sampling location and were thus considered local (Table S1 in Appendix S1). The remaining 33 individuals (26 males and seven females) were assigned to 10 genetically distinct populations 25 to 161 km from the spawning area and were classified as dispersers (Table S1 in Appendix S1).

Teno river salmon show significant variation in sea age at maturity, where spending more years at sea results in larger body size and higher reproductive success (Heinimaa & Heinimaa 2004; Reid & Chaput 2012). A range of sea age at maturity classes, and therefore sizes, were observed in both local and disperser fish (Table S4 in Appendix S1). As is common in many A tlantic salmon populations, the average sea age at maturity of females was higher than males (2.4 vs. 1.2 SW, respectively). Local and dispersers did not differ in sea age at maturity (females: 2.5 ± 0.1 SW, t = 1.55, P = 0.132, effect size 0.5 ± 0.3; males: 1.3 ± 0.1 SW, t = 0.98, P = 0.326, effect size 0.1 ± 0.1) or weight within sex (females: 8.06 ± 0.63 kg, t = 1.01, P = 0.322, effect size 0.58 ± 0.57; males: 3.81 ± 0.24 kg, t = 0.64, P = 0.525, effect size 0.17 ± 0.27; Table S4 in Appendix S1).

### Reproductive success

Bayesian parentage analysis assigned 1987 of the 5223 offspring (38%) to at least one sampled adult with confidence (Table S3 in Appendix S1). On average, local females were assigned 9.6 times more offspring than dispersing females (32.5 vs 3.4 offspring, respectively) and local males were assigned 2.9 times more than dispersing males (6.6 vs 2.3 offspring, respectively; Table 1, Fig. 2). These results were significant for both sexes (females, effect of origin as main effect: 1.38 ± 0.22, z = 6.18, p < 0.001; males, effect of origin: 0.54 ± 0.14, z = 3.75, p < 0.001; all models include main effects of sea age and number of sampled individuals, Table 1). This pattern of higher reproductive success among local females and males remained significant when restricting analyses to only include those adults that successfully reproduced at the site, i.e. only individuals positively assigned as parents to offspring (breeding individual reproductive success model: see Appendix S1 and Table S5 for further details).

**Table 1.**
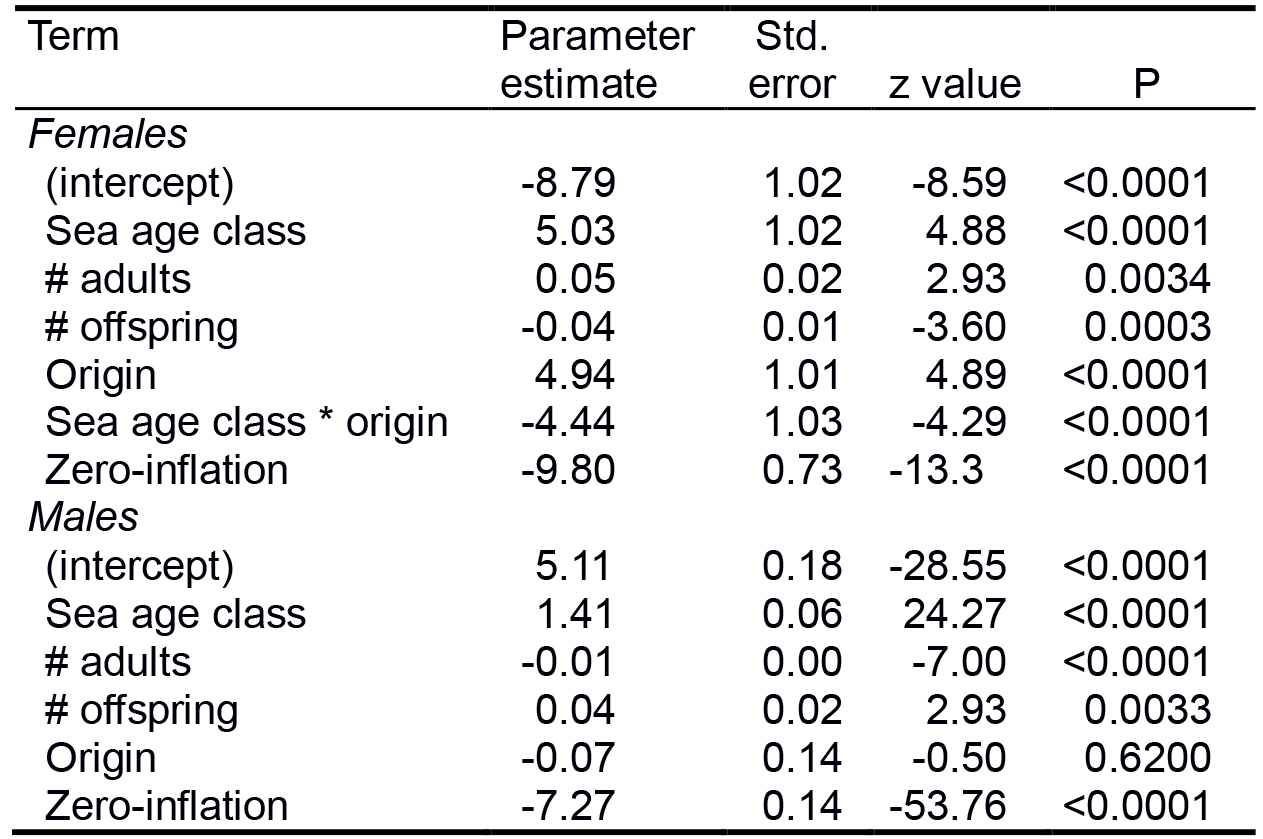
Summaries of minimal models testing the simultaneous effect on reproductive success of sea age at maturity as a two-level factor(age class), annual adult sample size(# adults) and offspring (# offspring), origin (local or disperser) and an sea age class * origin interaction(significant in females but removed in males because it was not significant)in adult males and females sampled at the lower Utsjoki location. The“zero-inflation” term gives the fitted intercept of the binomial component(whether a count was zero) of the mixture model. Each block of values shows the fit of one model.

| Term | Parameter estimate | Std. error | z value | P |
| --- | --- | --- | --- | --- |
| **Females** |  |  |  |  |
| (intercept) | -8.79 | 1.02 | -8.59 | <0.0001 |
| Sea age class | 5.03 | 1.02 | 4.88 | <0.0001 |
| # adults | 0.05 | 0.02 | 2.93 | 0.0034 |
| # offspring | -0.04 | 0.01 | -3.60 | 0.0003 |
| Origin | 4.94 | 1.01 | 4.89 | <0.0001 |
| Sea age class * origin | -4.44 | 1.03 | -4.29 | <0.0001 |
| Zero-inflation | -9.80 | 0.73 | -13.3 | <0.0001 |
| **Males** |  |  |  |  |
| (intercept) | 5.11 | 0.18 | -28.55 | <0.0001 |
| Sea age class | 1.41 | 0.06 | 24.27 | <0.0001 |
| # adults | -0.01 | 0.00 | -7.00 | <0.0001 |
| # offspring | 0.04 | 0.02 | 2.93 | 0.0033 |
| Origin | -0.07 | 0.14 | -0.50 | 0.6200 |
| Zero-inflation | -7.27 | 0.14 | -53.76 | <0.0001 |

The advantage of local spawners compared to dispersers was observed across all sizes and sea age maturation classes. Both males and females showed greater reproductive success with increasing sea age at maturity (males: 0.83 ± 0.03, z = 32.4, p < 0.001; females: 0.61 ± 0.05, z = 13.1, p < 0.001). In females, the difference in reproductive success between locals and dispersers was less pronounced in later maturing females (sea age class and origin interaction in addition to main effects of sea age, origin and number of adults and juveniles: −0.44 ± 1.03, z = −4.29, p < 0.001, Table 1, Fig. 2). Males, on the other hand, showed no evidence of different effects of origin in different sea age classes with local males being consistently more successful within each sea age maturation size class (sea age class and origin interaction: 0.13 ± 0.27, z = 0.492, p = 0.623; Table 1, Fig. 2).

### Consistency across cohort years

In both sexes, the fitness advantage of local spawners was robust across all cohort years, and remained significant when any single cohort year was removed from the dataset (i.e., on all three year subsets of the data with males and females pooled, the smallest effect of origin as a main effect on number of offspring was 1.79 ± 0.23 and all p < 0.001).

### Mating success and assortative mating

We examined the mating success (number of mates per individual) and whether mating was assortative according to origin (locals or dispersers). Among the 28 adult females and 115 males with at least one mate identified including unsampled mates, males had fewer mates than females (mean of 1.4 vs. 3.0 mates across both locals and dispersers, Table S4 in Appendix S1; effect of sex on number of mates as single main effect in Poisson GLM: −0.74 ± 0.13 log(mates), z = −5.54, p < 0.001). Among the 55 pairs in which both mates were identified, the majority consisted of two local individuals, whereas no successful disperser-disperser pairs were identified (50 pairs local-local, five mixed, four of which had a local mother). Hence, there was no indication of assortative mating with respect to origin, with local and dispersing parents having similar proportions of dispersing mates (effect sizes as logit(proportion of mates disperser): in males 14.7 ± 3269, z = 0.005, p = 0.996; in females, 15.5 ± 3097, z = 0.005, p = 0.996). Local and dispersing parents also did not differ in the weight of their mates (effect of origin on sea age at maturity-controlled weight of mates in males: 2.6 ± 0.6 kg, t = 1.06, p = 0.291; in females: 0.2 ± 0.4 kg, t = 0.61, p = 0.550) nor in the number of mates they had, although males had fewer mates than females overall (in a single model of number of mates: effect of origin 0.20 ± 0.30 log(mates), z = 0.69, p = 0.490; effect of sex −0.74 ± 0.13 log(mates), z = −5.57, p < 0.001). Despite the small number of mixed-origin pairs, they displayed a visibly lower reproductive success than local-local pairs (−0.85 ± 0.35, z = −2.45, p = 0.014).

## Discussion

It has been suggested that disperser disadvantage influences local adaptation by selecting against dispersal of adults and their offspring into new environments (Hendry 2004; Nosil *et al.* 2005). Our aim in this study was to utilize natural straying events to assess the occurrence of selection against dispersal in the wild. Although the majority of adults sampled had indeed returned to their natal spawning grounds, there was nevertheless considerable potential for gene flow between genetically distinct populations with dispersing adults accounting for approximately 12.5% of the potential parental pool. Despite this, the home ground advantage of locals resulted in several times higher reproductive success for both males and females over dispersers. We also found an increase in reproductive success with sea age at maturity, indicating that the older, and therefore larger, fish tend to have higher reproductive potential, regardless of origin.

Dispersal and migration likely play important roles in phenotypic trait divergence (Kinnison *et al.* 2003; Edelaar & Bolnick 2012). However, we found no phenotypic differences in size and sea age at maturity between local and dispersing individuals. Costs of dispersal in terms of physical migration to new locations, arrival precedence on the spawning grounds, and familiarity with the natal environment may give local breeders an advantage, yet these factors are difficult to measure in the wild (Kinnison *et al.* 2003; Peterson *et al.* 2014). On the other hand, sex-specific and juvenile straying may promote adult dispersal (Hamann & Kennedy 2012), but evidence for factors that cause these behavioral differences is limited (Quinn 1993; Hendry *et al.* 2003; Peterson *et al.* 2016; Bett *et al.* 2017). If phenotypic differences between local and dispersing individuals were observed (e.g., Peterson *et al.* 2014) then assortative mating by phenotype may reinforce genetic and phenotypic differentiation. Our examination of mate choice, albeit constrained by the low number of mating pairs including dispersers, gave no clear evidence of assortative mating based on origin, be it local or disperser. This result implies that mating preferences for local mates is sufficiently low to allow dispersers to obtain mates in non-natal localities.

An alternative explanation for our observations could be that some adults caught at the site did not actually attempt to spawn in this location, but instead continued their migration to tributaries or the mainstem further upriver. In our study, dispersers almost exclusively originated from localities further upstream of our study location and thus it is possible they were still migrating upstream to spawn. Studies in salmonids show that movements in male sockeye salmon (*Oncorhynchus nerka*) decreases as the spawning season progresses (Rich *et al.* 2006), and that most dispersers first homed to their natal streams before dispersing to new spawning grounds (Peterson *et al.* 2016). Studies of Atlantic salmon caught in the Teno river show that movement of individuals to other areas is rare close to spawning (Økland *et al.* 2001) and radiotelemetry of individuals has also shown very limited movement during the breeding season (P Karppinen & J Erkinaro, unpublished data, see also Karppinen & Erkinaro 2009). Thus, long distance movement of individuals visiting this spawning site close to spawning time is highly unlikely. Finally, in the unlikely event that dispersers were to spawn in several locations within a year, their reproductive success on non-natal spawning grounds remains demonstratively lower than local spawners, even when accounting for only individuals that spawned. Therefore, a reproductive disadvantage need not be universal for the individual disperser, but specific to the location such that dispersers underperform when spawning on a non-natal habitat but potentially still produce more offspring, and thus obtain higher fitness, in their natal habitat.

### Local adaptation

A fitness dispersal disadvantage raises the strong possibility that local adaptation will evolve if this pattern is observed throughout the system. A similar assessment of reproductive success in a single cohort at a second location in the Teno system (Akujoki, see Appendix S1) allowed us to partially test for local adaptation. Mirroring our main results from lower Utsjoki, local individuals from Akujoki also had higher reproductive success over dispersers (local vs. dispersing females, 11.7 vs. 8.2 offspring; local vs. dispersing males, 9.0 vs 1.6 offspring; see Appendix S1, Fig. S2, Tables S1, S3-4, S6-7). However, this is effect is only significant in males, possibly due to low sample sizes (Fig. S2 and Tables S4 in Appendix S1)

A key advantage of the current study compared to traditional reciprocal transplantation experiments is that we assessed reproductive success of natural dispersal events. However, our natural dispersal approach also precluded a fully reciprocal design, and hence unambiguous identification of local adaptation. We have demonstrated a dispersal disadvantage in terms of reproductive success, yet the factors responsible for this pattern remain unclear as we found no phenotypic differences between locals and dispersers in either study location. Thus, the potential for local adaptation is high in spite of appreciable adult dispersal, although the ecological factors driving such local adaptation remain unresolved in this case. Using genomic screening, a recent study identified river flow volume as a potentially adaptively important environmental characteristic in Teno River Atlantic salmon (Pritchard *et al.* 2018). Future research on ecological differences between localities may reveal whether habitat preferences drive local adaptation.

## Conclusion

This study provides convincing empirical support for fine-scale local selection against dispersal in a large Atlantic salmon meta-population, signifying that local individuals have a marked home-ground advantage in reproductive success. These results emphasize the notion that migration and dispersal may not be beneficial in all contexts and highlighting the potential for selection against dispersal and for cryptic local adaptation to drive population divergence across fine geographic scales.

## Supporting information

## Acknowledgements

We thank Katja Salminen, Meri Lindqvist, Jenni Kuismin, Jani Aaltonen, Susanna Ukonaho and Jan Laine for laboratory assistance, Jorma Kuusela, Jari Haantie and Matti Kylmäaho for scale aging analyses, and Olavi Guttorm, Topi Pöyhönen, Timo Kanniainen, Arto Koskinen, Jorma Ollila, Mari Lajunen, Tuomo Karjalainen (deceased), Mikko Kytökorpi, Seda Karslioglu, Anna Ellmen and Hans Pieski for field assistance.

## Author contributions

C.R.P., J.E., P.O., T.A. & ME designed the study. M.E., J.-P.V., P.O., S.J. & C.R.P. collected samples. M.E. & J.-P.V. performed laboratory work. H.G.-W. & J.-P.V. analysed the data, K.B.M., C.R.P. & H.G.-W. wrote the manuscript, with input and final approval from all other authors.

## Ethics

Fishing permission for research purposes was granted by the Lapland Centre for Economic Development, Transport, and the Environment (permit numbers 1579/5713-2007 and 2370/5713-2012).

## Appendix S1

Supplementary materials and methods, results, Figures S1-2, Tables S1-7, supplementary references.

