## Supplementary material for "Home ground advantage: selection against dispersers promotes local adaptation in wild Atlantic salmon"

**Appendix S1:** Supplementary materials and methods, results, Figures S1-2, Tables S1–7, supplementary references.

### Home ground advantage: selection against dispersers promotes local adaptation in wild Atlantic salmon

Kenyon B. Mobley[1†*], Hanna Granroth-Wilding[1,2†], Mikko Ellmen[2], Juha-Pekka Vähä[3], Tutku Aykanat[1], Susan E. Johnston[4], Panu Orell[5], Jaakko Erkinaro[5], Craig R. Primmer [1,6,7]

[1]*Organismal and Evolutionary Biology Research Program, Faculty of Biological and Environmental Sciences, PO Box 56, 00014 University of Helsinki, Finland*
[2]*Department of Biology, University of Turku, Finland, Itäinen 10 Pitkäkatu 4, Turku FI-20520, Finland*
[3]*Association for Water and Environment of Western Uusimaa, POB 51, FI-08101 Lohja, Finland*
[4]*Institute of Evolutionary Biology, University of Edinburgh, Edinburgh EH9 3FL, UK*
[5]*Natural Resources Institute Finland (Luke), PO Box 413, FI-90014 Oulu, Finland*
[6]*Insititue for Biotechnology, 00014 University of Helsinki, Finland*
[7]*Helsinki Institute of Sustainability Science, 00014, University of Helsinki, Finland*

†Shared first authorship

#### Supplementary Materials and Methods

##### Teno River

The Teno River supports one of the world's largest and most phenotypically diverse Atlantic salmon stocks (Niemelä *et al.* 2006; Erkinaro *et al.* 2018). Up to 50,000 individuals are harvested by local fishermen and recreational fisheries annually (Erkinaro *et al.* 2018), representing up to 20% of the riverine Atlantic salmon catches in Europe (ICES 2011). The Utsjoki river is one of the largest tributaries of the Teno river system (length 66 km, catchment area 1652 km$^2$), draining into the main stem 108 km from the Barents sea.

##### Lower Utsjoki sampling location

Our focal sampling location covered the first kilometres from the mouth of the Utsjoki tributary, referred to hereafter as lower Utsjoki (Fig. 1, main text). The lower Utsjoki harbors several distinctive spawning grounds (c. 150–400 m) which are separated by 150–600 m river sections with pools and slow flowing reaches. Wetted widths of the spawning areas vary from 30-50 m and the maximum spawning site depth is ~300 cm, although most nests (redds) are at depths between 70-150cm. Thermal conditions in lower Utsjoki are strongly influenced by Lake Mantojärvi (194 ha, max depth 60 m), situated approximately 5 km upstream from the river mouth: the lowest river stretch freezes over usually 1–2 weeks later than in the Teno main stem or in other tributaries. The daily mean water temperatures typically drops below 7$^o$C by the end of September and the peak spawning activity usually takes place during the first week of October.

The lower Utsjoki location has been the focus of numerous research projects and monitoring programmes (Pihlaja *et al.* 1998; Niemelä *et al.* 1999; Niemelä *et al.* 2001; Mäki-Petäys *et al.* 2002)) and includes permanent monitoring sites where annual electrofishing surveys have been conducted since 1979 (Niemelä *et al.* 1999). Therefore, it is highly unlikely that areas with substantial numbers of juvenile salmon would be overlooked in our juvenile sampling. Regions with both high and low juvenile density were sampled each year with a view to ensuring the sampling of offspring produced by individuals spawning in different quality habitats.

Repeat spawning

A total of 13 individuals (6 female, 7 male) that had spawned in a previous year were identified based on scale morphology (repeat spawners, or kelts) were captured at the lower Utsjoki location (4.9% of all adult individuals captured in this location). The mean sea age at maturity of repeat spawning females was 3.2 ± 0.4 SE (range2-4 SW) and all repeat spawning males had spent one year at sea before the first spawning migration and another year at sea before returning to spawning for the second time (all repeat spawners were 2SW). Only one repeat spawning female was a disperser, all other individuals were local. Additionally, adult genotypes were screened for recaptures using allelematch (Galpern *et al.* 2012). Allelematch identifies full and partial genotype matches based on genotype data. Using the criterion of up to two allele mismatches, we found no recaptured adults pooled across all locations (lower Utsjoki and Akujoki) and cohort years (2010-2014).

##### Breeding versus non-breeding individuals in reproductive success models

Reproductive success was modelled in a mixture model that allowed instances of zero reproductive success to be considered both due to parents not attempting to breed at all and due to a spawning attempt being made but failing to produce any offspring surviving to, or detected at, sampling. To investigate whether dispersal disadvantage remains significant among breeding individuals, we repeated our main analyses on a dataset including only those adults assigned as parents to offspring excluding individuals that did not mate. However, using only individuals that mated did not change the main conclusions of the study. Using the reproductive success of the 115 males and 28 females that had offspring assigned to them as parents, we tested for a main effect of origin alongside main effects of sea age at maturity and annual sample size of adults and offspring as in the main analysis. As these data were not zero-inflated, we used generalized linear models with Poisson errors; effect sizes were log-transformed, as fitted by the model. Among both males and females that produced offspring, local individuals had higher reproductive success, with a more pronounced effect in females, just as in the full analysis, with local males producing on average 9.8 offspring compared to 7.7 for dispersers, and local females 30.9 compared to 12.0 for dispersers (effect of origin on number of offspring: in males, $0.51 \pm 0.14$, $z = 3.70$, $p = 0.001$; in females, $1.33 \pm 0.21$, $z = 6.37$, $p < 0.001$).

**Selection against dispersers at a second sampling location, Akujoki**

To assess whether selection against dispersers is a general feature of the river system, a parent-offspring cohort from 2011-2012 from the Akujoki tributary of the Teno River was analysed. Sampling, parentage analysis, population assignment and statistical methods were similar to those conducted for lower Utsjoki.

**Akujoki sampling location**

The Akujoki river (69°35'5.82"N, 25°57'46.73"E) is a small tributary of the Teno river with a drainage area of 193 $km^2$ located 80km upstream from the lower Utsjoki location (Fig. 1 maintext). It. It flows c. 35 km through a mountain valley before connecting to the River Teno main stem 192 km upstream from the sea. Only the lower 6 km of the river is accessible by salmon due to an impassable waterfall. Wetted widths of the spawning areas range between 10 - 30 m and the maximum depth is <1 m. The peak spawning season in the Akujoki is between mid-September and the first week of October (Orell *et al.* 2011). Adults were sampled during the $2^{nd}$ week of September 2011 using gill nets and offspring were collected by electrofishing in September 2012.

**Population assignment**

Fishes from the Akujoki tributary are genetically distinct from the Teno mainstem (Vähä *et al.* 2017) (Table S6 in Appendix S1). The baseline sample from the Akujoki tributary location was significantly differentiated from all other sampling locations (Vähä *et al.* 2017) (Table S6 in Appendix S1).

A total of 63 adults (32 males and 31 females) and 607 juveniles were sampled at the Akujoki spawning grounds (Table S1 in Appendix S1). The adult sex ratio of the Akujoki location did not differ from equality (two-sided $\chi^2 = 0.01$, $P = 0.920$). The Akujoki population contained a significantly greater proportion of disperser fish than the Utsjoki population (two-sided $\chi^2 = 48.41$, $P < 0.0001$) with 41.3% of adults hailing from the local population and the remainder from adjacent populations (Table S1 in Appendix S1). Sixteen of 32 males (50.0%) were considered local while only 10 of the 31 females were considered local (32.3%).

**Parentage assignment**

At the Akujoki locality, 94 offspring with fewer than 7 loci successfully genotyped were excluded, leaving a total of 513 offspring in the analysis; all 63 adults were successfully genotyped for at least 11 of the 13 loci (46 at all 13 loci). 97.9% of offspring were confidently assigned at least one parent including unsampled adults and 73.7% of offspring (378 individuals) were confidently assigned to at least one sampled adult (Table S3 in Appendix S1).

**Adult phenotypic characteristics**

Local and dispersers were phenotypically similar to one another, mirroring results from lower Utsjoki (weight: females $t = -0.23$, $P = 0.814$, effect size -0.03 ± 0.15; males $t = 0.17$, $P = 0.868$, effect size 0.02 ± 0.11, Table S3). Females also did not differ in sea age between local and dispersers ($t = 0.36$, $P = 0.724$, effect size = 0.1 ± 0.2; all males were 1 SW Table S4 in Appendix S1). In Akujoki, both females and males were younger and hence lighter than in lower Utsjoki (females, mean sea age 1.3 SW (range 1-2SW), mean weight 1.9 kg; males, all aged 1SW, mean weight 1.4 kg Table S4 in Appendix S1). All sampled males at the Akujoki location matured after one year at sea and therefore sea age affects could not be tested.

**Reproductive success**

Reproductive success in the Akujoki location was calculated and analysed using similar approaches as for lower Utsjoki with the following exceptions. As sampling in Akujoki was carried out in only one year, annual sample size was not included in these models. The sea age at maturity of all Akujoki males was one sea winter, so sea age at maturity was not included as a predictor in the male analysis. Interactions of origin with sea age at maturity could not be tested in Akujoki females due to the very low number of females maturing after two or more sea winters.

In Akujoki, local individuals had higher reproductive success than dispersing individuals (local vs. dispersing females, 11.7 vs. 8.2 offspring; local vs. dispersing males, 9.0 vs 1.6 offspring, respectively, Fig. S2 and Tables S4 in Appendix S1), but this effect of origin was only significant in males (males, effect of origin on number of offspring: 1.95 ± 0.22, $z = 8.76$, $p < 0.001$; females, effect of origin: -0.02 ± 0.12, $z = -0.13$, $p = 0.896$, Fig. S2 and Table S7 in Appendix S1). These results were very sensitive to two high-fecundity outliers with strong statistical leverage: a disperser 2SW female, 4.3 kg, with 100 offspring and a local 1SW male, 1.8 kg, with 109 offspring. These two individuals were assigned together as a parent pair to 11 offspring. When these outlier parents were excluded, local fish of both sexes had higher reproductive fitness: local females had 3.2 times more offspring than dispersing females (means of 11.7 and 3.7 offspring, respectively) and local males 1.4 times more (mean 2.3 and 1.6 offspring, respectively). The effect of origin on number of offspring, in males excluding outlier, 0.755 ± 0.27, $z = 2.84$, $p < 0.005$; in females excluding outlier, 0.891 ± 0.17, $z = 5.39$, $p < 0.001$.

**Repeat spawning**

A slightly larger proportion of individuals (7.9%) captured at Akujoki were repeat spawners as compared to the lower Utsjoki location. A total of 5 repeat spawning females, all with a sea-age of 2SW, were identified based on scale morphology. Two of these repeat spawning females were foreign while the remaining three females were from Akujoki.

**Figure S1**. Interpolation of sea age at maturity from weight for sampled female (F) and male (M) adult salmon for which scales could not be sampled. The pale histograms show the distribution of weights for individuals of known sea age and the thick lines show calculated normal distributions with the mean and standard deviation of the samples. For ease of interpretation, calculated distributions are scaled to the count data. Points jittered around the x-axis show the weights of the sampled individuals of unknown sea age. These were assigned an sea age based on the calculated distribution under which that weight was most likely to occur and are colored according to their assigned sea age. Triangles show local individuals and circles show disperser individuals. Note differences in scale between males and females.

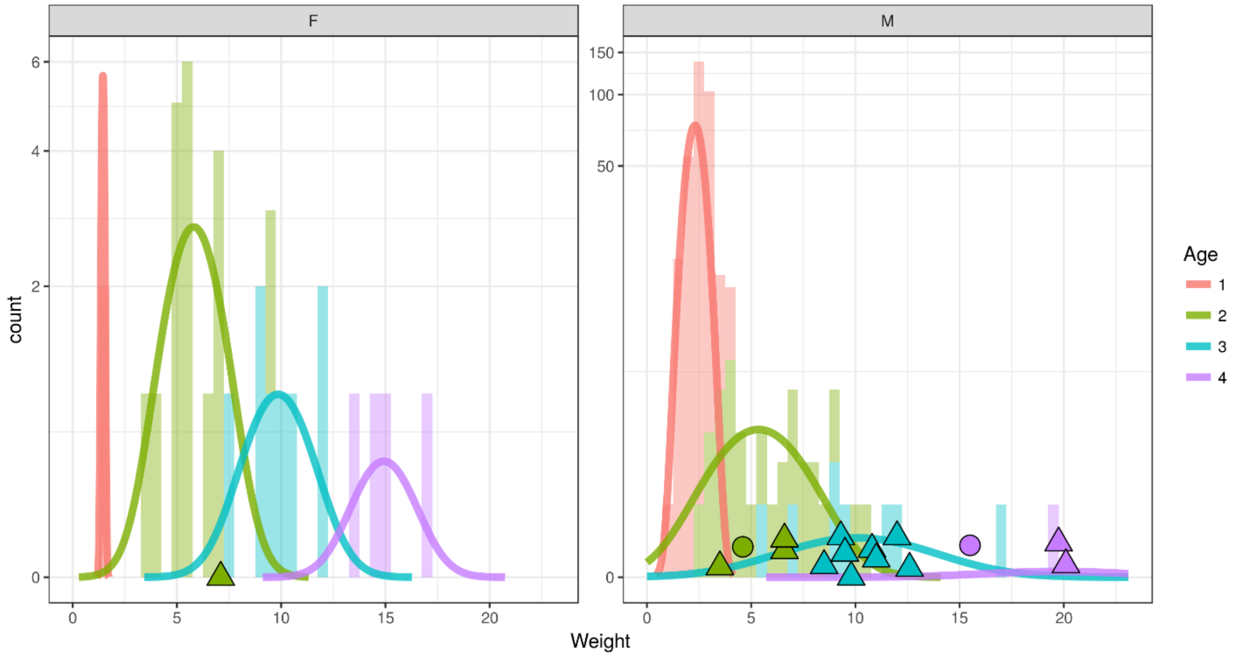

**Figure S2.** The effect of individual origin on reproductive success in females and males. The Akujoki tributary. Small symbols show the number of offspring produced by each individual, and the large symbol with error bars the mean ± one standard error of the mean for each combination of sea age at maturity and origin. For clarity, points are jittered on the x axis and the y-axis is shown on a log scale. In females, the effect of origin was masked by one highly fecund (100 offspring) disperser (point shown).

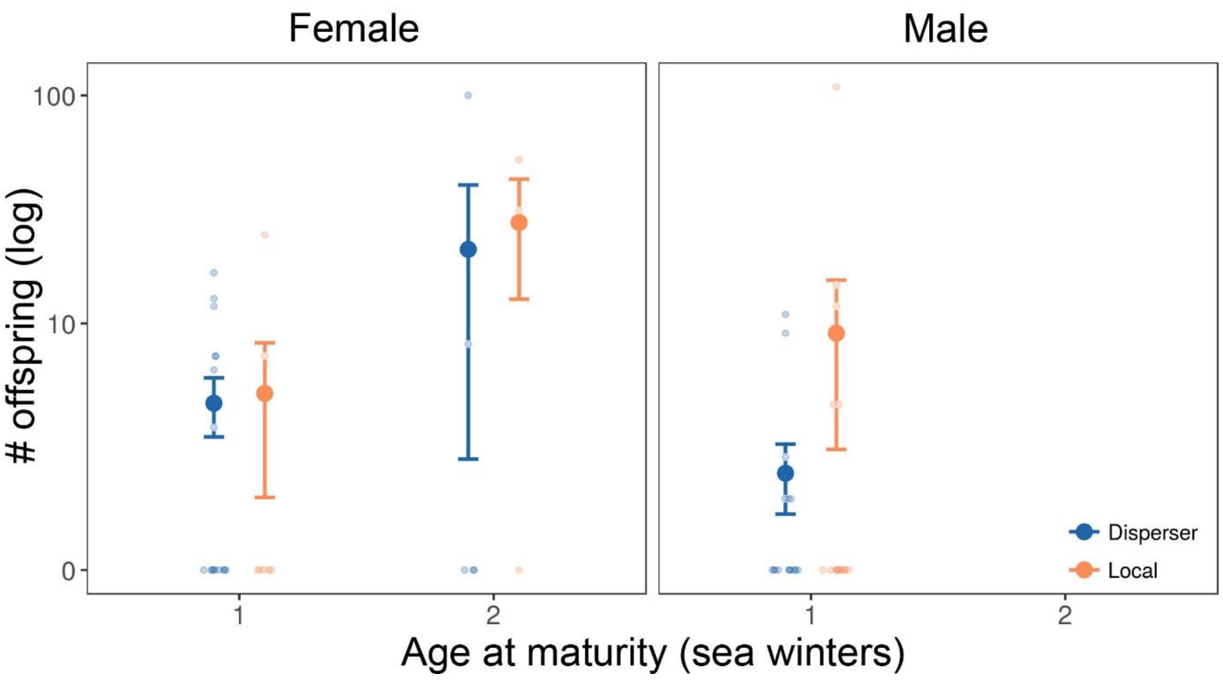

**Table S1.** Baseline population adult assignments for lower Utsjoki sampling location. Origin, abbreviation (abb. in Fig. 1 main text), number of adults assigned to the baseline population (males (M), females (F) and total) and overwater distance from lower Utsjoki (distance, taken from Vähä *et al.* (2017))

| Origin | Abb. | M | F | total | Distance (km) |
|---|---|---|---|---|---|
| Local | | | | | |
| Lower Utsjoki | UL | 205 | 27 | 232 | -- |
| Foreign | | | | | |
| Upper Utsjoki | UU | 4 | 3 | 7 | 8.0 |
| Tsarsjoki | TZ | 0 | 1 | 1 | 25.0 |
| Kevojoki | KE | 1 | 1 | 2 | 25.0 |
| Vetsijoki | VJ | 1 | 0 | 1 | 37.0 |
| Yläköngäs | YK | 3 | 5 | 8 | 50.5 |
| Levajohka | LJ | 1 | 0 | 1 | 55.0 |
| Outakoski | OU | 8 | 0 | 8 | 103.5 |
| Akujoki | AJ | 2 | 0 | 2 | 109.5 |
| Goššjohka | GS | 3 | 0 | 3 | 170.0 |
| Inari | IN | 2 | 0 | 2 | 183.0 |
| Total | | 25 | 10 | 35 | |

Table S2. Adult sample sizes in each cohort and sampling location. Unsampled population sizes were estimated as part of the pedigree fitting using MasterBayes.

| | Lower Utsjoki | | | | Akujoki |
|---|---|---|---|---|---|
| | 2011 | 2012 | 2013 | 2014 | 2011 |
| Offspring | 821 | 1008 | 1523 | 1871 | 607 |
| Adults | 54 | 34 | 66 | 110 | 63 |
| Males (local/disperser) | 44/3 | 23/4 | 43/11 | 94/8 | 16/16 |
| Females (local/disperser) | 7/0 | 5/2 | 8/4 | 7/1 | 10/21 |
| Estimated unsampled parent population | | | | | |
| Males (median) | 287.7 | 196.9 | 158.9 | 198.5 | 56.3 |
| 95% quantiles | 243.7-341.3 | 168.4-231.2 | 142.8-177.4 | 181.0-218.2 | 46.9-67.7 |
| Females (median) | 31.9 | 23.5 | 37.8 | 72.0 | 18.8 |
| 95% quantiles | 26.7-38.4 | 20.3-27.3 | 33.7-42.7 | 61.9-84.0 | 15.6-22.6 |
| % male pop. captured | 14.0 | 12.1 | 25.4 | 33.9 | 36.2 |
| % female pop. captured | 18.0 | 23.0 | 24.1 | 10.0 | 62.2 |

**Table S3.** Parentage assignments using the MasterBayes pedigree framework. "Confidently unsampled parent" refers to assignments in which all sampled adults can be confidently (> 90% likelihood) excluded as potential parents. “No confident assignment” refers to cases where no single genotyped adult could be confidently assigned as a parent, nor could all genotyped parents be confidently excluded. Our analyses were based on assignments where at least one parent was confidently positively identified, i.e. where the natal population of at least one parent was known.

| | Lower Utsjoki | | | | | Akujoki |
|---|---|---|---|---|---|---|
| | 2011 | 2012 | 2013 | 2014 | Total | 2011 |
| Two sampled adults confidently assigned as parents | 39 | 44 | 119 | 105 | 307 | 81 |
| One sampled adult confidently assigned, the other confidently unsampled | 215 | 319 | 554 | 592 | 1680 | 282 |
| % offspring with at least one sampled adult confidently assigned as a parent† | 31.8 | 36.3 | 44.5 | 37.5 | | 73.7 |
| Total offspring genotyped | 821 | 1008 | 1523 | 1871 | 5223 | 513 |

†In cases where only one parent was identified among the sampled adults, either the other parent is assigned as a confidently unsampled individual or no confident assignment of the other parent could be made.

**Table S4.** Mean sea age at maturity (sea age in seawinters, SW), weight in kg (weight), mating success (# mates) and reproductive success (# offspring) of local and dispersing adults of each sex in the lower Utsjoki and Akujoki locations. The table shows the number of individuals (n) and mean values (± standard error of the mean) for each group. Data from lower Utsjoki are pooled over cohort years. In Akujoki, all males matured after one year at sea and thus sea age effects were not tested.

| Population | Sex | Origin | n | Sea Age (SW) | Weight (kg) | # mates | # offspring |
|---|---|---|---|---|---|---|---|
| Lower Utsjoki | Female | Local | 27 | 2.5 ± 0.1 | 8.06 ± 0.63 | 3.1 ± 0.4 | 32.5 ±7.7 |
| Lower Utsjoki | Female | Disperser | 7 | 2.0 ± 0.4 | 5.38 ± 1.68 | 1.5 ± 0.5 | 3.4 ± 3.3 |
| Lower Utsjoki | Male | Local | 204 | 1.3 ± 0.0 | 3.81 ± 0.24 | 1.4 ± 0.1 | 6.6 ± 1.1 |
| Lower Utsjoki | Male | Disperser | 26 | 1.2 ± 0.1 | 3.23 ± 0.59 | 1.6 ± 0.3 | 2.3 ± 1.2 |
| Akujoki | Female | Local | 10 | 1.3 ± 0.2 | 1.94 ± 0.40 | 1.8 ± 0.5 | 11.7 ± 5.9 |
| Akujoki | Female | Disperser | 21 | 1.2 ± 0.1 | 1.82 ± 0.25 | 2.0 ± 0.4 | 8.2 ± 4.7 |
| Akujoki | Male | Local | 16 | - | 1.38 ± 0.08 | 2.2 ± 0.8 | 9.0 ± 6.8 |
| Akujoki | Male | Disperser | 16 | - | 1.36 ± 0.08 | 1.3 ± 0.3 | 1.6 ± 0.8 |

**Table S5.** Summaries of generalized linear models (GLM) of reproductive success only using males and females known to have bred. Models test the effect on reproductive success of sea age at maturity as a two-level factor (sea age class), annual adult sample size (# adults) and offspring (# offspring), origin (local or disperser) and an sea age class * origin interaction (removed because it was not significant) in adult males and females sampled at the lower Utsjoki location. GLMs were fitted with Poisson errors; effect sizes were log-transformed, as fitted by the model Each block of values shows the fit of one model.

| Term | Parameter estimate | Std. error | z value | P value |
|---|---|---|---|---|
| *Females* | | | | |
| (intercept) | -6.50 | 0.30 | -21.75 | < 0.001 |
| Sea age class | 0.60 | 0.05 | 13.11 | < 0.001 |
| # adults | 0.02 | 0.02 | 1.32 | 0.188 |
| # offspring | -0.02 | 0.01 | -1.69 | 0.091 |
| Origin | 1.33 | 0.21 | 6.37 | < 0.001 |
| *Males* | | | | |
| (intercept) | -6.66 | 0.19 | -35.03 | < 0.001 |
| Sea age class | 0.82 | 0.03 | 32.68 | < 0.001 |
| # adults | -0.01 | 0.00 | -7.39 | < 0.001 |
| # offspring | 0.06 | 0.01 | 3.99 | < 0.001 |
| Origin | 0.51 | 0.14 | 3.70 | < 0.001 |

**Table S6.** Baseline population adult assignments for the Akujoki sampling location. Natal location, abbreviation (abb. in Fig. 1 main text), number of adults assigned to the baseline population (males (M), females (F) and total) and overwater distance (Vähä *et al.* 2017) from Akujoki.

| Origin | Abb. | M | F | total | Distance (km) |
|---|---|---|---|---|---|
| Local | | | | | |
| Akujoki | AJ | 16 | 10 | 26 | 0.0 |
| Disperser | | | | | |
| Outakoski | OU | 2 | 2 | 4 | 6.0 |
| Nilijoki | NJ | 8 | 10 | 18 | 34.5 |
| Báišjohka | BJ | 3 | 2 | 5 | 35.5 |
| Karigasjoki | KR | 2 | 4 | 6 | 35.5 |
| Iskurasjoki | IJ | 1 | 1 | 2 | 56.5 |
| Kuoppilasjoki | KJ | 0 | 1 | 1 | 73.0 |
| Lower Teno River† | AK, PI, SI, GJ, KO | 1 | 0 | 1 | 109.5 |
| Total | | 33 | 30 | 63 | |

†Distance taken from the average from these locations (Vähä *et al.* 2017)

**Table S7.** Model summaries showing the effect of age at maturity as a two-level factor (sea age class, 1 or 2SW) and origin (local or disperser) on reproductive success of males and females sampled in Akujoki. Among both males and females there was one outlier that had a strong influence on model fits (see Appendix SI materials and methods and Akujoki results for details); the upper model set shows models with the outliers retained and the lower set shows models with these high-leverage outliers removed. Male models did not include sea age class as all sampled males spent only one year at sea. The "zero-inflation" term gives the fitted intercept of the binomial component (whether a count was zero) of the mixture model.

| Model and terms | Parameter estimate | Std. error | z value | p |
|---|---|---|---|---|
| **All data** | | | | |
| *Females* | | | | |
| (intercept) | -5.36 | 0.22 | -24.87 | < 0.001 |
| Sea age class | 1.50 | 0.13 | 11.64 | < 0.001 |
| Origin | -0.02 | 0.12 | -0.13 | 0.896 |
| Zero-inflation | -5.20 | 0.36 | -16.25 | < 0.001 |
| *Males* | | | | |
| (intercept) | -4.83 | 0.21 | -23.42 | < 0.001 |
| Origin | 1-95 | 0.22 | 8.76 | < 0.001 |
| Zero-inflation | -9.61 | 0.37 | -14.99 | < 0.001 |
| **Excluding high-leverage outliers** | | | | |
| *Females* | | | | |
| (intercept) | -4.80 | 0.23 | -21.05 | < 0.001 |
| Sea age class | 0.66 | 0.16 | 4.11 | <0.001 |
| Origin | 0.89 | 0.17 | 5.39 | <0.001 |
| Zero-inflation | -5.84 | 0.37 | -15.66 | < 0.001 |
| *Males* | | | | |
| (intercept) | -4.83 | 0.21 | -23.48 | < 0.001 |
| Origin | 0.75 | 0.27 | 2.84 | 0.005 |
| Zero-inflation | -5.51 | 0.39 | -14.28 | < 0.001 |
